# Transcriptomic comparison of in vitro models of the human placenta

**DOI:** 10.1101/2024.06.14.598695

**Authors:** Samantha Lapehn, Sidharth Nair, Evan J. Firsick, James MacDonald, Ciara Thoreson, James A Litch, Nicole R. Bush, Leena Kadam, Sylvie Girard, Leslie Myatt, Bhagwat Prasad, Sheela Sathyanarayana, Alison G. Paquette

## Abstract

Studying the human placenta through in vitro cell culture methods is necessary due to limited access and amenability of human placental tissue to certain experimental methods as well as distinct anatomical and physiological differences between animal and human placentas. Selecting an in vitro culture model of the human placenta is challenging due to representation of different trophoblast cell types with distinct biological roles and limited comparative studies that define key characteristics of these models. Therefore, the aim of this research was to create a comprehensive transcriptomic comparison of common in vitro models of the human placenta compared to bulk placental tissue from the CANDLE and GAPPS cohorts (N=1083). We performed differential gene expression analysis on publicly available RNA sequencing data from 6 common in vitro models of the human placenta (HTR-8/SVneo, BeWo, JEG-3, JAR, Primary Trophoblasts, and Villous Explants) and compared to CANDLE and GAPPS bulk placental tissue or cytotrophoblast, syncytiotrophoblast, and extravillous trophoblast cell types derived from bulk placental tissue. All in vitro placental models had a substantial number of differentially expressed genes (DEGs, FDR<0.01) compared to the CANDLE and GAPPS placentas (Average DEGs=10,873), and the individual trophoblast cell types (Average DEGs=5,346), indicating that there are vast differences in gene expression compared to bulk and cell-type specific human placental tissue. Hierarchical clustering identified 53 gene clusters with distinct expression profiles across placental models, with 22 clusters enriched for specific KEGG pathways, 7 clusters enriched for high-expression placental genes, and 7 clusters enriched for absorption, distribution, metabolism, and excretion genes. In vitro placental models were classified by fetal sex based on expression of Y-chromosome genes that identified HTR-8/SVneo cells as being of female origin, while JEG-3, JAR, and BeWo cells are of male origin. Overall, none of the models were a close approximation of the transcriptome of bulk human placental tissue, highlighting the challenges with model selection. To enable researchers to select appropriate models, we have compiled data on differential gene expression, clustering, and fetal sex into an accessible web application: <u>“Comparative Transcriptomic Placental Model Atlas (CTPMA)”</u> which can be utilized by researchers to make informed decisions about their selection of in vitro placental models.

## Introduction

The placenta is the first fetal organ that develops during pregnancy, and is involved in many processes crucial for fetal growth and development and parturition, including supplying oxygen and nutrients to the fetus [1]. The placenta develops from the trophectoderm, which is the outer layer of the preimplantation embryo, or blastocyst [2]. The trophectoderm mediates adhesion of the blastocyst to the uterine epithelium which leads to stable contact of the embryonic and maternal tissues, leading to placenta formation [3]. The placenta is composed of cells including fibroblasts, immune and vascular cells, and tissue-specific trophoblast cells that are responsible for the placenta’s unique functions [2]. The trophoblast differentiates into extravillous trophoblasts, cytotrophoblasts, and syncytiotrophoblasts which each maintain unique roles in placental physiology [2,3]. Extravillous trophoblasts are invasive cells that anchor the placenta to the maternal decidua and play a role in remodeling of the spiral arteries to increase blood flow and nutrient transport to the fetus [4,5]. Cytotrophoblast and syncytiotrophoblast cells make up placental villous trees, with syncytiotrophoblasts forming the outer layer while cytotrophoblasts form the layer beneath where they serve as progenitor cells to expand and replenish the syncytiotrophoblast layer. As the outer layer of the villous trees, syncytiotrophoblasts interface with maternal blood and are thus the primary exchange surface of the placenta for maternal-fetal transfer of oxygen, nutrients, hormones, and waste [1].

There are numerous in vitro models of the human placenta from primary and immortalized sources that represent the various trophoblast cell types of the placenta. HTR-8/SVneo is an immortalized cell line of extravillous trophoblast cells collected from first trimester placentas [6] and represents a heterogeneous population of trophoblast and stromal cells [7]. Cells collected from choriocarcinomas, which are highly malignant human chorionic gonadotrophin (hCG) secreting tumors that can originate from gestational and non-gestational tissues, are comprised of trophoblast cells and can also be used to model the placenta [8]. Placental cell lines derived from gestational choriocarcinomas include JEG-3, JAR and BeWo. BeWo can model the transition from cytotrophoblasts to syncytiotrophoblasts, as these cells fuse together with the addition of forskolin, an activator of cyclic AMP [9]. JEG-3 and JAR cells are also choriocarcinoma cells that are representative of the cytotrophoblast, but do not syncytialize in culture [10].

There are also in vitro models derived directly from human placental tissue. Indeed, primary trophoblast cells which are collected and cultured from placental tissue and can spontaneously syncytialize in culture [11] over a period of 3 days, recapitulating in vivo syncytiotrophoblast formation. Placental villous explants are another primary tissue model, which consist of excised tissue samples of placental villous trees that are cultured in vitro [12]. Being derived from primary tissue, these trophoblast cells and placental villous explants can be used to understand placental physiology and the prenatal environment in the context of maternal and fetal health characteristics, since demographic data, medical outcomes, and prenatal exposures can be collected for these samples.

To date, there have been several comparative studies of in vitro placental models, but many are limited to a specific physiological parameter or to a small number of cell lines without direct comparison to human tissue. Examples of this work include: an evaluation of BeWo, JAR, JEG-3, and HTR-8/SVneo cells as a suitable model for preeclampsia drug screening [13], an assessment of BeWo, JAR, JEG-3, and ACH-3P as models for endocrine and transport studies [10], and DNA methylation profiling of JAR, BeWo, JEG-3, HTR-8/SVneo, AC1M-32, TEV-1, and SWAN71 [14]. Additional studies include an analysis of HTR-8/SVneo, JEG-3, JAR and BeWo for epithelial and mesenchymal cell markers [7], studies of invasive properties of HTR-8/SVneo and JEG-3 cells [15,16], a quantitative proteomic evaluation of xenobiotic and steroid metabolizing enzymes in placental tissue and BeWo, JEG-3, JAR, and HTR-8/SVneo cell lines [17], as well as studies of single genes or small gene subsets across multiple placental cell lines [18,19]. Overall, these studies reveal that in vitro models of the human placenta have individual strengths that should be considered during model selection; however, they have not directly compared these models to human placental tissue. RNA sequencing data is well-suited to characterize these in vitro placental models because it gives a comprehensive overview of gene expression which can be extrapolated to understand differences in physiological parameters and protein function. RNA sequencing data is also easily available through online data repositories such as Gene Expression Omnibus (GEO) and Sequence Read Archive (SRA) with increasing frequency due to open science initiatives.

The use of placental cell models is an important tool since human placental samples are difficult to obtain and not amendable to some types of experimental manipulation. Animal models of the placenta are also not ideal as they are highly species-specific and not a good reflection of human anatomy and physiology [20,21]. However, there is limited knowledge about how closely these cell and primary tissue models recapitulate the expression of genes and their corresponding biological pathway networks in bulk placental tissue. The overall goal of this project was to use publicly available RNA sequencing data from placental cell lines to determine which placental cell model was closest to human placental tissue. This project summarizes the transcriptomic landscape of common placental cell lines and primary tissue models to provide a comprehensive, gene level summary of similarity to bulk placental tissue and individual trophoblast cell types with the results made broadly accessible through the development of an interactive web application.

## Materials and Methods

### Placental Tissue Samples from CANDLE and GAPPS

RNA sequencing data of placental tissue samples was derived from 2 cohorts: the Conditions Affecting Neurocognitive Development and Learning in Early Childhood (CANDLE) study and Global Alliance to Prevent Prematurity and Stillbirth (GAPPS) study by ECHO PATHWAYs [22]. CANDLE enrolled participants with uncomplicated singleton pregnancies between the ages of 16-40 years that delivered at participating hospitals in Shelby County, TN. CANDLE placenta collection has been previously described and included dissection of a rectangular piece of placental villous tissue (∼2cm x 0.5cm x 0.5cm) from the middle of the parenchyma within 15 minutes of delivery. Tissue samples were refrigerated overnight in RNAlater at 4°C before storage at -80°C in RNAlater . GAPPS enrolled pregnant participants that were 18 years and older (or medically emancipated) that were recruited in Seattle, WA and Yakima, WA. GAPPS placenta collection has been previously described [23,24] and included four 8mm vertical tissue punches from the placental disc that were collected from 2 sites (separated by 7cm). Tissue samples were placed in RNAlater within 30 minutes of delivery and stored at - 80°C. Tissue collected as part of both studies were manually dissected and cleared of maternal decidua. Use of these samples for this study was approved by the University of Washington Institutional Review Board (IRB). Research activities related to the CANDLE study were approved by the University of Tennessee Health Sciences Center IRB while research activities related to the GAPPS samples was approved by the IRB of the Seattle Children’s Research Institute. ECHO PATHWAYs generated RNA sequencing data on 794 samples from CANDLE and 289 samples from GAPPS, as previously described [22] (Supplemental Figure 1).

### Download of Publicly Available Data

Publicly available RNA sequencing data of select in vitro models of the human placenta were identified through a search of cell model names on GEO. Data were downloaded from GEO using the GEOquery R package, confirmed to contain unfiltered count data, and then filtered to include only control samples [25]. Treatments of control samples included non-targeting knockdowns (i.e. siRNA, shRNA), vehicle controls (i.e. DMSO), and untreated samples. Data that were not available as unfiltered counts on GEO were downloaded from the NCBI Sequence Read Archive (SRA) using sratools. Transcript abundances were estimated from the downloaded SRA fastq files using the pseudo-alignment program kallisto, with bias correction using genome alignment GRCh38 [26]. Transcripts were then condensed to Ensembl Gene IDs using tximport producing count data [27]. A summary of the datasets used in this analysis is available with GEO accession numbers and citations in Table 1 and is summarized in Supplemental Figure 1.

**Table 1:** Control treated RNA sequencing data of in vitro placental models including Villous Explants, Primary Trophoblasts, JAR cells, JEG-3 cells, BeWo cells, and HTR-8/SVneo cells were downloaded from SRA or GEO. Placental tissue data was acquired from the CANDLE and GAPPS cohorts and are available on dbGaP (CANDLE (N=794): phs003619.v1.p1, GAPPS (N=298): phs003620.v1.p1).

| Placental Model | GEO Accession and Citation | Number of Control Samples | Syncytialized? |
| --- | --- | --- | --- |
| **Primary Trophoblasts** | GSE245394 [59] | 6 | Yes |
|  | GSE217210 [30] | 14 | Yes |
|  | Paquette Lab Data | 16 | Yes |
| **HTR-8/SVneo** | GSE10305 [60] | 3 | NA |
|  | GSE217210 [30] | 6 | NA |
|  | GSE197810 [61] | 3 | NA |
|  | GSE154339 [40] | 8 | NA |
|  | GSE85995 [62] | 3 | NA |
| **BeWo** | GSE66962 [63] | 3 | Yes |
|  | GSE197810 [61] | 3 | No |
|  | GSE200983 | 3 | No |
|  | GSE171648 [64] | 3 | No |
|  | GSE106394 | 2 | No |
|  | GSE72437 | 5 | No |
|  | GSE175815 [65] | 6 | Some Samples |
|  | GSE209620 [66] | 3 | No |
|  | Paquette Lab Data | 15 | Some Samples |
| **JEG-3** | GSE143859 [67] | 4 | NA |
|  | GSE185071 [68] | 2 | NA |
|  | GSE72437 | 5 | NA |
|  | GSE106394 | 2 | NA |
|  | GSE192446 [69] | 3 | NA |
| **Villous Explants** | GSE154489 [40] | 5 | NA |
|  | GSE104348 [41] | 3 | NA |
| **JAR** | GSE106394 | 2 | NA |
|  | GSE216043 [70] | 3 | NA |
|  | GSE192446 [69] | 3 | NA |
|  | GSE199278 [71] | 5 | NA |
| **CTBs** | GSE182381 [39] | 3 | No |
| **EVTs** | GSE182381 [39] | 5 | NA |
| **STBs** | GSE182381 [39] | 3 | Yes |

### Data Processing and Merging

All datasets were merged by Ensembl gene IDs to include the union of all genes included in individual datasets (N= 20,052 genes). We then filtered the data to include only protein-coding genes. Genes that were undetected (NA) in individual datasets were set to 0 counts, then the data was transformed to log counts per million (Supplemental Figure 1). Data was filtered to remove genes that had an average logcpm<0 across all samples, which is in line with our previous treatment of low-expressing genes [28–30]. Our final dataset included 12,793 genes. We normalized data using the trimmed mean of M values method [31]. To correct for samples originating from different labs and sequencing platforms, we used the “remove unwanted variation using residuals” (RUVr) function from the RUVSeq R package [32] to account for unknown confounding and batch effects. The weights calculated from RUVr were added to the design matrix for differential gene expression analysis. Data variability was assessed and visualized through principal components analysis.

### Differential Gene Expression Analysis

Differential gene expression analysis was conducted using the limma-voom pipeline from the limma R package [33]. Linear models were generated to compare the placental tissue data from CANDLE and GAPPS to the BeWo, JEG-3, JAR, Primary Trophoblasts, HTR-8/SVneo, and Villous Explants, termed the “Placental Tissue Models”. Additional linear models were generated to compare the BeWo (syncytialized and unsyncytialized), HTR-8/SVneo, JAR, JEG-3, and primary trophoblasts to single trophoblast cell types (extravillous trophoblasts (EVTs), cytotrophoblasts (CTBs), and syncytiotrophoblasts (STBs)). EVTs and CTBs were derived using fluorescence-activated cell sorting (FACS) whereas STBs were derived from villous tissue digests. The results of this analysis will be referred to as the “Trophoblast Cell Type RNAseq Models”. P-values were adjusted for multiple hypothesis testing using the Benjamini-Hochberg method and genes with FDR<0.01 were considered significant [34].

### Hierarchical Clustering Analysis

Gene expression data (N=12,793) was averaged across samples by placental model then used to perform hierarchical clustering using average linkage and Euclidean distance. Clusters of genes with similar expression profiles across the different placental models were combined using the R package dynamicTreeCut [35] with a tree cut height of 8 and a minimum cluster size of 50 genes. Pathway analysis for each gene cluster was performed using the limma R package function kegga [33] to run an overrepresentation enrichment test on KEGG pathways (excluding disease-related pathways). The 12,793 genes used to construct the hierarchical clusters that were mapped to at least 1 KEGG pathway were used as background. P-values were corrected for multiple hypothesis testing using the Benjamini-Hochberg approach and pathways with FDR<0.05 were considered significant.

### Genes of Interest Evaluation

Genes with elevated gene expression in the placenta were downloaded from the Human Protein Atlas (HPA), selecting genes classified as placenta specific (N=11), placenta enriched (N=54; 4x higher expression compared to any other tissue), or placenta enhanced (N=169; 4x higher expression compared to the average of the other tissues) [36].

In vitro models are frequently used for performing experimental treatments that cannot be studied in vivo. It is therefore important that the model selected for performing in vitro treatments expresses the appropriate receptors and metabolic pathways for in vivo comparison. To evaluate the potential for placental models to recapitulate the drug metabolizing and enzymatic functions of placental tissue, a list of absorption, distribution, metabolism, and excretion genes from the PharmaADME consortium were downloaded [37]. Each of these lists of genes was evaluated for average expression by placental model, differential gene expression, and cluster membership. After assigning cluster membership, HPA and ADME genes were evaluated for statistical overrepresentation within each overlapping cluster using a one-sided Fisher’s Exact Test (p<0.05).

### Fetal Sex Determination

Unfiltered count data from each HTR-8/SVneo, BeWo, JAR, and JEG-3 dataset and each individual villous explant and primary trophoblast cell sample was processed as follows to quantify gene expression on the Y chromosome. Count data was transformed to counts per million and Y chromosome expression was summarized as both total counts per million of the Y chromosome and average counts per million per Y chromosome gene. Samples with total counts per million less than 10 for the Y chromosome were classified as female samples whereas samples with total counts per million greater than 265 were classified as male samples [38].

## Results

In this analysis, we integrated publicly available data from 21 datasets across 6 in vitro models of the human placenta, as well as 1083 human placental tissue samples from 2 birth cohorts to evaluate the suitability of various cell lines to recapitulate placental gene expression data. First, we performed Principal Component Analysis (PCA) to evaluate the overall variance in the data (Figure 1). For the placental tissue models, the first three PCs accounted for 44.8% of the variance (Figure 1A), while the trophoblast cell type RNAseq models, the first three PCs account for 50.1% (CTBs, Figure 1B), 49.6% (EVTs, Figure 1C), and 50.6% (STBs, Figure 1D) of the variance. For all four models, clustering was largely based on in vitro model type. In the placental tissue models, the CANDLE/GAPPS placental tissue samples were most closely clustered with the primary trophoblasts, HTR-8/SVneo, and BeWo cells based on PC1 (Figure 1A). For all three of the trophoblast cell types (EVTs, CTBs, STBs), primary trophoblast cells were most closely clustered (Figure 1B, 1C, 1D). Across all 4 PCA plots, JEG-3 and JAR cells which are both derived from choriocarcinomas are closely clustered (Figures 1A, 1B, 1C, 1D). Separating the BeWo cells by syncytialization status for comparison to the villous tissue digest STBs showed distinct separation of syncytialized and unsyncytialized BeWo, although neither showed close clustering with the STBs for each PCA (Figure 1B, 1C, 1D).

**Figure 1:**
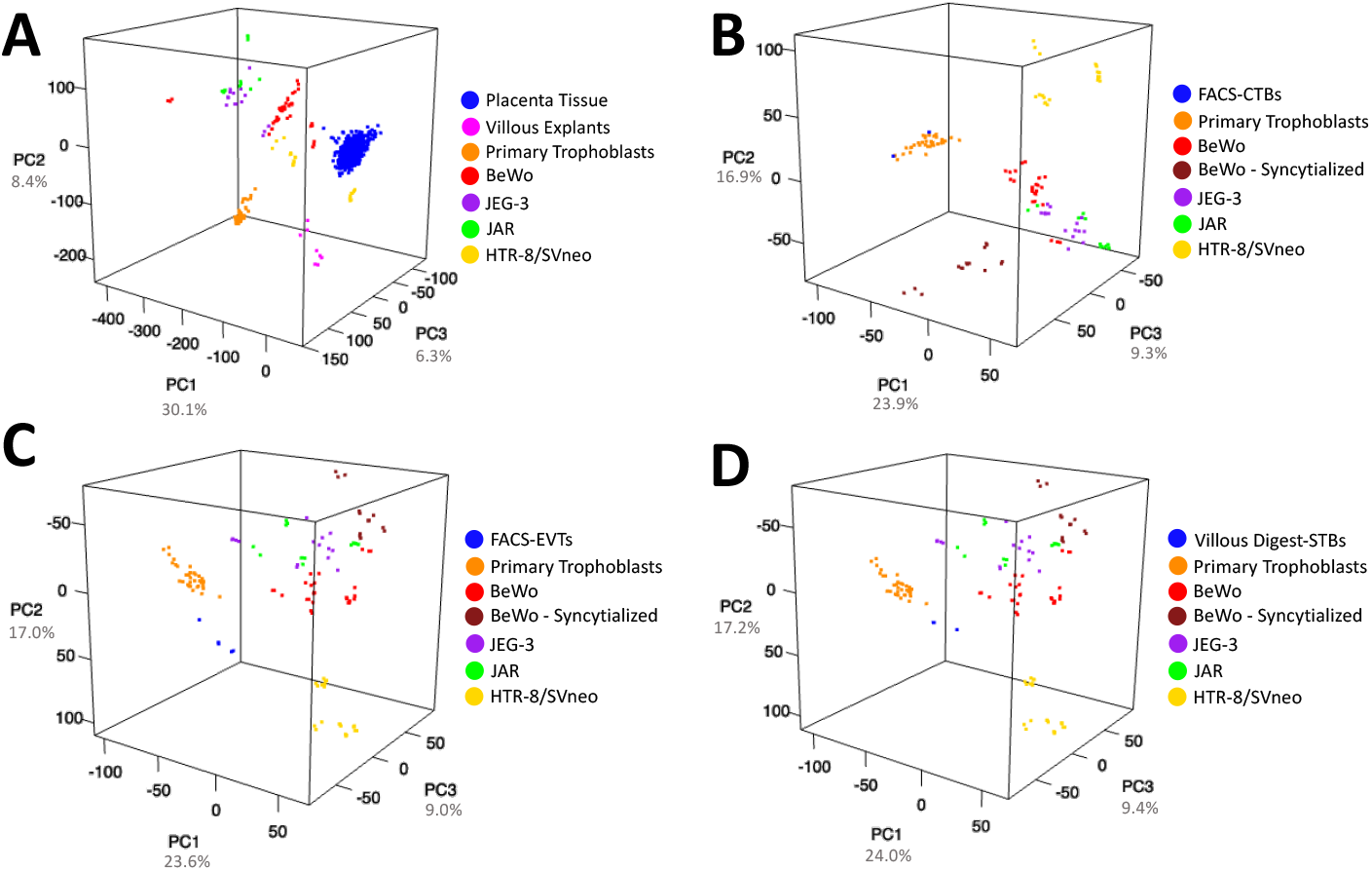
Principal Components Analysis (PCA) for the in vitro placental models compared to A) CANDLE and GAPPS placental tissue, B) FACs Cytotrophoblasts (CTBs), C) FACS Extravillous Trophoblasts (EVTs), and D) Villous tissue digest Syncytiotrophoblasts (STBs). Variance explained by each of the first three PCs is indicated in the axis labels.

Next, we evaluated transcriptomic differences at the gene level using a series of linear regression models. In the placental tissue model, which compared gene expression of in vitro placental models to gene expression from CANDLE and GAPPS placentas, the villous explant samples had the lowest number of DEGs (9,885), while the HTR-8/SVneo cells had the largest number of DEGs (11,639) (Figure 2A). Across all three trophoblast cell type RNAseq models (CTBs, EVTs, STBs), the models comparing primary trophoblasts to these trophoblast cell types had the lowest number of DEGs (CTBs-4291, EVTs-5416, STBs-4275), while JEG-3 had the highest number of DEGs (CTBs-5804, EVTs-6696, STBs-5635) (Figure 2B, 2C, 2D). The results of this analysis provide evidence that the cell-based models of the placenta are closer in gene expression to these primary tissue cell sorted samples than the bulk placental tissue, but still with substantial gene expression differences (Average DEGs=5,346). Full results of the differential gene expression analysis are available in Supplemental Table 1 (Placental Tissue Models) and Supplemental Table 2 (Trophoblast Cell Type RNAseq Models-CTBs, EVTs, STBs). In summary, all the evaluated in vitro models of the human placenta exhibit substantial gene expression differences compared to human bulk placental tissue or individual trophoblast cell types derived from primary tissue indicating that selection of in vitro models should be driven by biological processes or functions of the placenta relevant to specific research questions.

**Figure 2:**
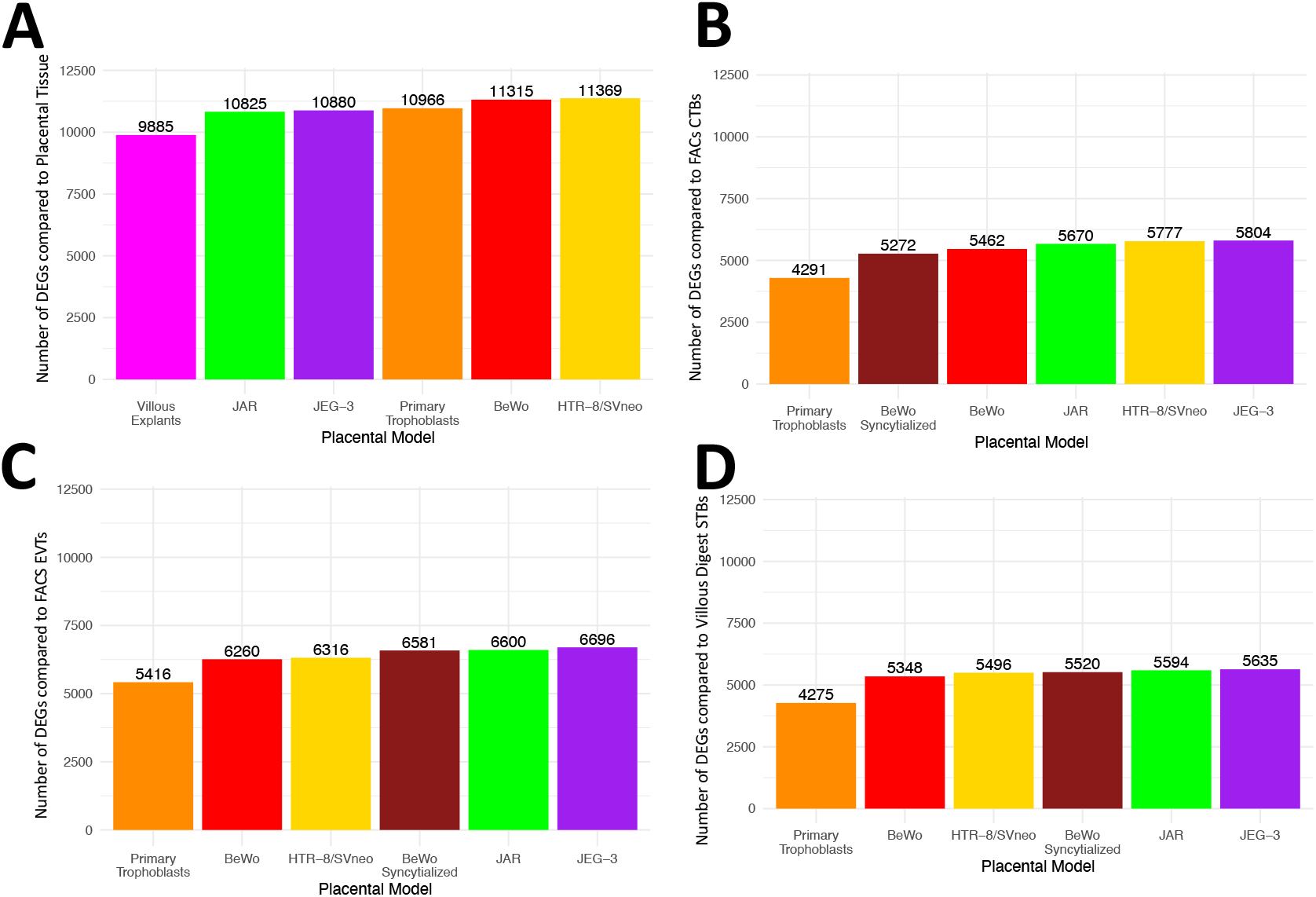
The number of differentially expressed genes (FDR<0.01) for each placental model after comparison to A) CANDLE and GAPPS placental tissue, B) FACs Cytotrophoblasts (CTBs), C) FACS Extravillous Trophoblasts (EVTs), and D) Villous tissue digest Syncytiotrophoblasts (STBs).

We sought to further define unique biological pathways and subgroups specific to individual placental models using hierarchical clustering and pathway enrichment analysis. Clustering of gene expression data from the placental tissue models (12,793 genes) generated 53 clusters of genes (Figure 3A). The largest cluster was Cluster 1 and contained 1353 genes, and the smallest cluster was Cluster 52 which contained the minimum allowed cluster size of 50 genes. The average cluster size was 241 genes. By averaging the expression of genes within each cluster by placental model, patterns of unique gene expression by placental model emerged (Figure 3B). For example, some clusters like 1, 2, 3, 9, and 32 exhibited similar expression profiles across all groups indicating that any of the in vitro models could be a good proxy for evaluating gene expression of one of these genes. However, other clusters such as 17 indicate that there is lower expression of this gene subset in the commercial choriocarcinoma cell lines (JEG-3, JAR, BeWo), which could be important if wanting to study one of these genes in vitro. Several clusters (Clusters 5, 11, 12, 23, 24, 34, 37, 38) show lower expression in Villous Explants compared to the rest of the groups.

**Figure 3:**
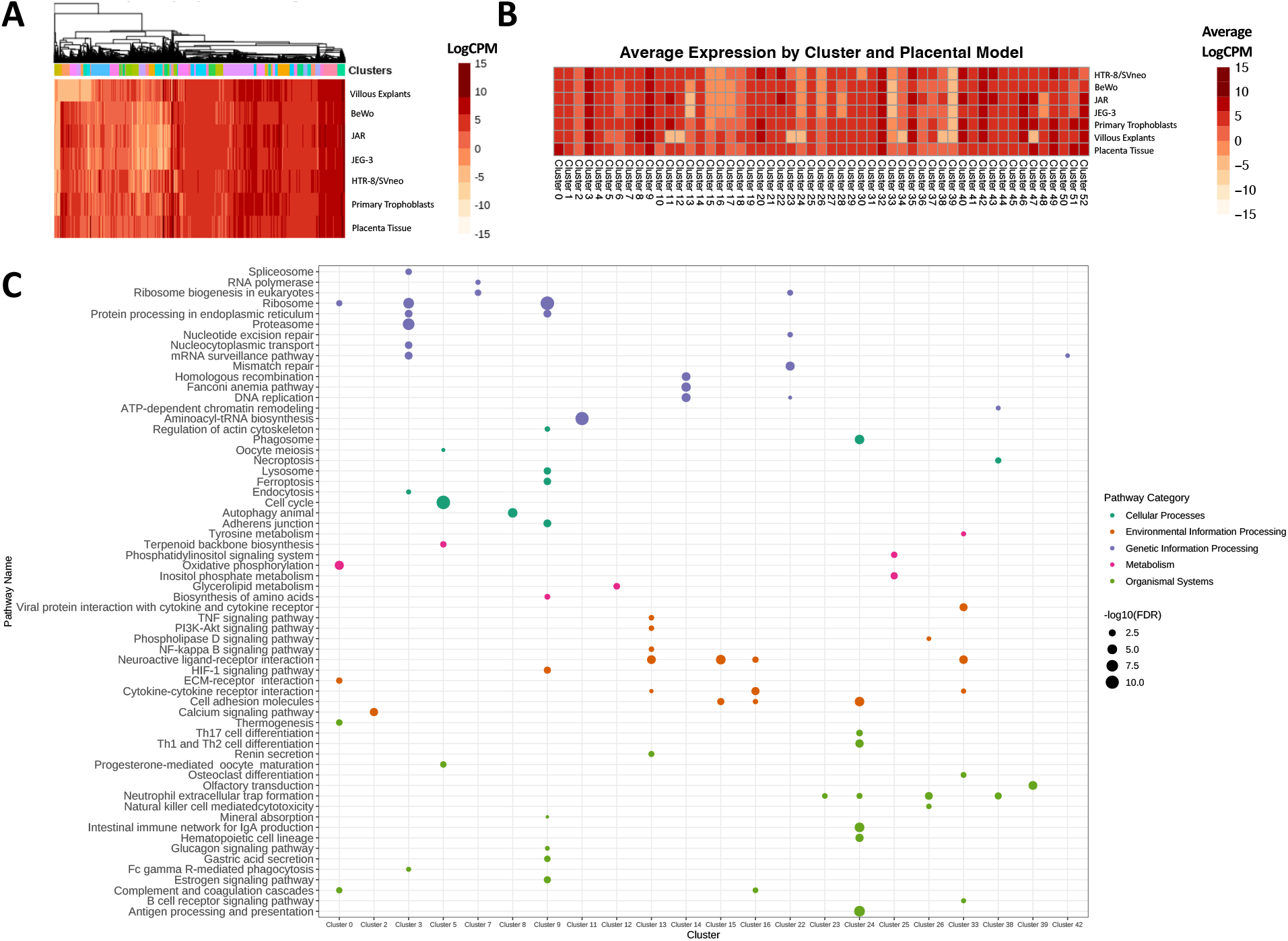
A) Hierarchical clustering of expression data from the CANDLE and GAPPS placental tissue model (N=12,793 genes) identified 53 clusters of genes. B) Average Expression of all genes within each cluster across each placental model. C) Significant KEGG pathways (FDR<0.05) enriched for genes that are members of individual clusters.

Pathway enrichment analysis was performed to gain insight into the biological functions of these clusters. 22 of 53 clusters were enriched for 1 or more KEGG pathways (FDR<0.05) (Figure 3C). 63 pathways were enriched for at least one cluster, with 51 of these pathways being unique to a single cluster (Figure 3C). Clusters that were enriched for single pathways included, Cluster 2-Calcium Signaling Pathway, Cluster 8-Autophagy, Cluster 11-Aminoacyl tRNA Biosynthesis, Cluster 12-Glycerolipid Metabolism, Cluster 23-Neutrophil Extracellular Trap Formation, Cluster 39-Olfactory Induction, and Cluster 42-mRNA Surveillance Pathway. Of note, Cluster 42 is one of the few clusters that shows lower expression in the bulk placental tissue compared to the in vitro models, while Clusters 12 and 23 show substantially lower expression in the Villous Explants. Overall, this pathway analysis provided insight into the biological functions of these clusters and indicates biological processes that may be different between these models.

Genes with elevated expression in the placenta compared to other human tissues as detailed by the Human Protein Atlas (HPA) were evaluated for cluster membership to see if distinct patterns of expression exist for these high-expression placental genes in placental tissue compared to in vitro placental models. We identified 8 placenta specific, 53 placenta enriched, and 157 placenta enhanced genes expressed in our data (Figure 4). Overall, most of these genes exhibited higher expression in the bulk placental tissue from CANDLE and GAPPS and the in vitro models derived from primary tissue sources (Primary Trophoblasts and Villous Explants) compared to the immortalized placental cell lines (Figure 4). There were 7 clusters (Clusters 0, 15, 16, 26, 30, 51, 52) that contained a statistical overrepresentation (p<0.05) of HPA gene (Supplemental Table 3). Of these overrepresented clusters, Clusters 0 (*PSG3*, *PSG11*, *PSG8*, *PSG9*), 16 (*LGALS13*), 30 (*ERVV*-2), and 51 (*PLAC1*) each contained 1 or more of the 8 placental HPA genes that are exclusively expressed in the placenta (Supplemental Table 1). While the genes in Cluster 0 show high expression across all models, the other 6 clusters overrepresented for placental HPA genes exhibit varying degrees of expression, with Cluster 51 lower in choriocarcinoma cell lines (JEG-3, BeWo, JAR) and Cluster 52 lower in HTR-8/SVneo cells compared to all other models which could be relevant to researchers using these placental model systems to study these placental genes.

**Figure 4:**
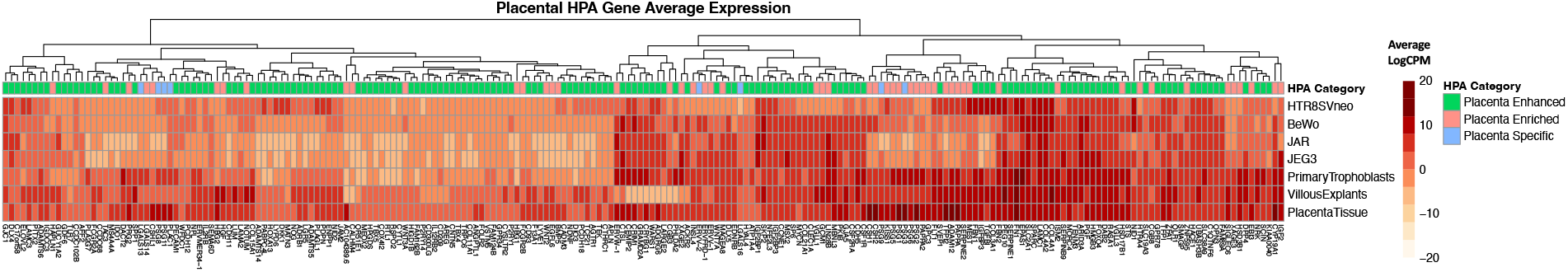
Average expression of genes of interest for placental biology including placenta specific, placenta enriched, and placenta enhanced genes defined by the Human Protein Atlas. 218 genes of 234 placental HPA genes were expressed at sufficient levels (average logcpm>0) in our placental models.

We similarly investigated the expression and cluster membership of ADME genes since in vitro placental models are often used for the study of pharmacological or toxicological agents that may depend on the expression of specific metabolic genes and processes. Of 298 ADME genes from PharmaADME, there were 158 expressed in the placental tissue model dataset including 11 core ADME genes and 147 extended ADME genes (Figure 5). Differential gene expression by placental model, HPA or ADME categorization, and hierarchical cluster assignments of these genes can be found in Supplemental Table 1. There were 7 other clusters (Clusters 2, 15, 16, 17, 28, 30, 51) that contained a statistical overrepresentation of ADME genes (Supplemental Table 3). Cluster 2 is overrepresented with 17 ADME genes (Supplemental Table 1) but based on average expression across the cluster (Figure 3B), there is stable expression across all placental models indicating that any of the tested models could be suitable for studying these genes. Conversely, Cluster 30 which is overrepresented with 5 ADME genes (*SLC7A7, NOS3, SLC22A11, ALDH3B2, SLC13A3*) (Supplemental Table 1) shows lower expression in HTR-8/SVneo cells (Figure 3B) indicating this model may not be adequate for experiments that require adequate expression of these genes and their associated biological pathways.

**Figure 5:**
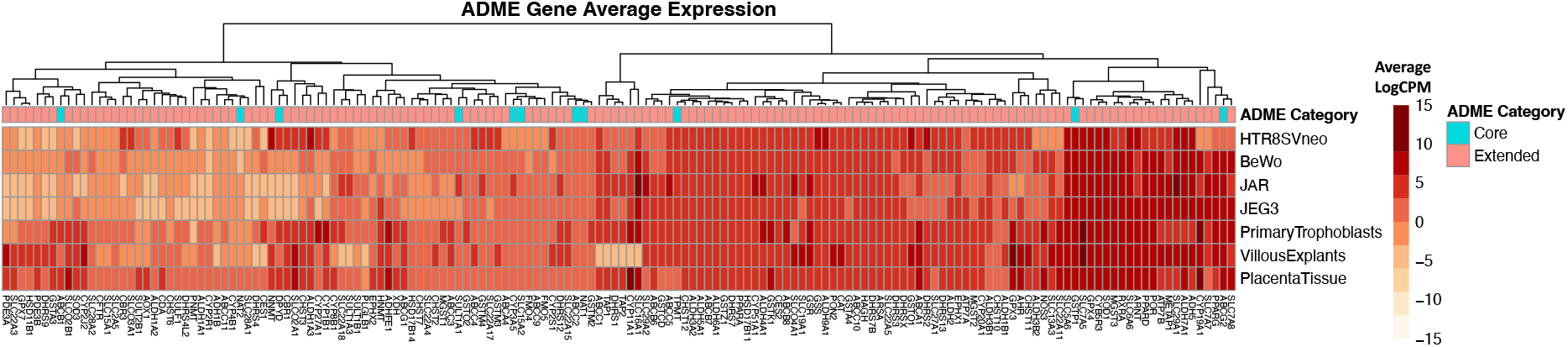
Average expression of Absorption, Distribution, Metabolism, and Excretion (ADME) genes from PharmaADME. 158 genes of 298 ADME genes were expressed at sufficient levels (average logcpm>0) in our placental models to be included.

Fetal sex of each placental model was determined based on quantifying the gene expression of the Y-chromosome. Based on this assessment, BeWo, JAR, and JEG-3 cell lines are of male origin, while the HTR-8/SVneo cell line is of female origin (Figure 6, Supplemental Table 4). As expected, the expression of the Y chromosome corresponds to the biological sex of the samples in the primary placental models (Primary Trophoblasts and Villous Explants) (Figure 6).

**Figure 6:**
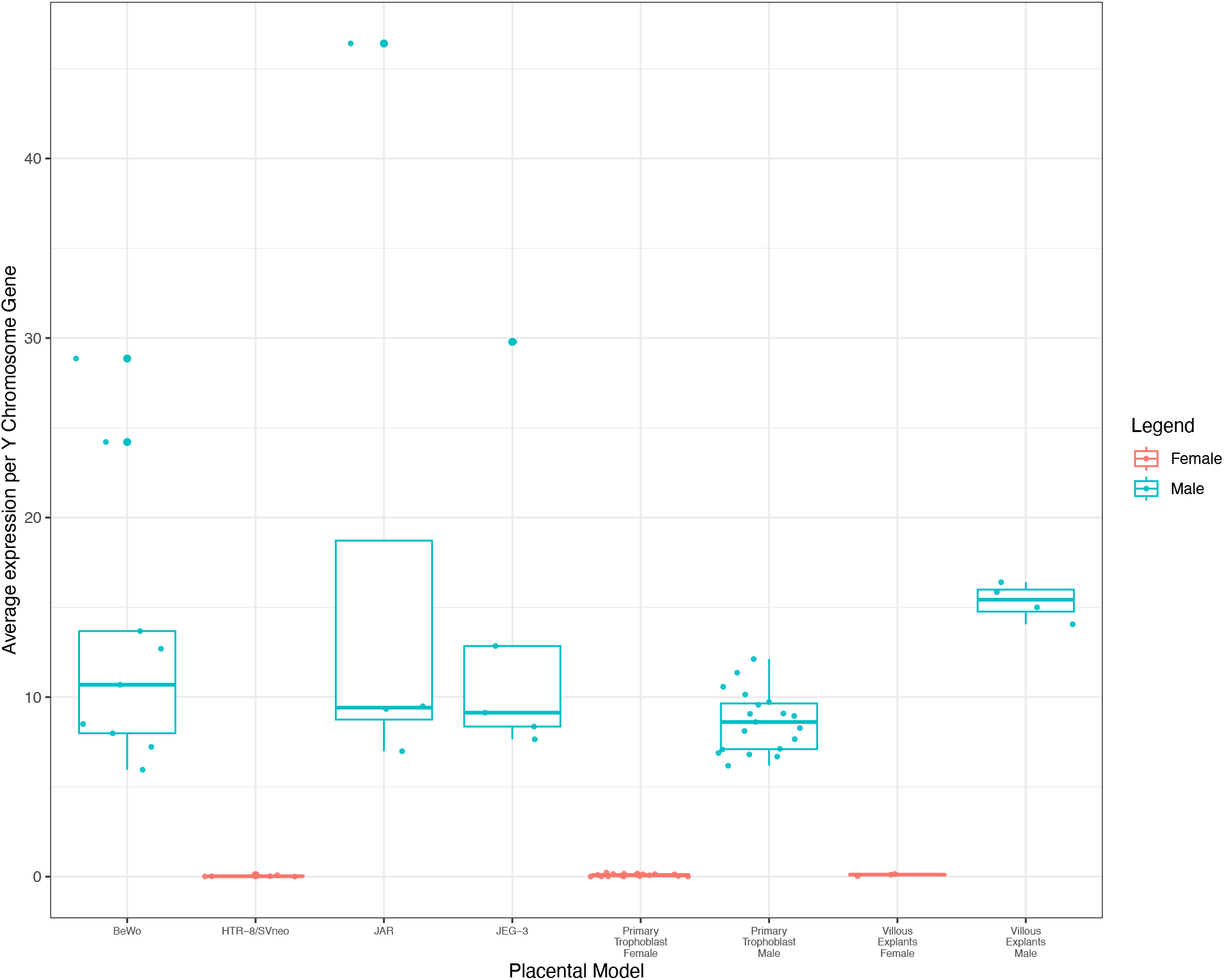
Fetal sex of placental models as determined by quantification of Y-chromosome gene expression. Placental models with mixed fetal sex (Villous Explants and Primary Trophoblasts) are displayed separately by sex.

Results of this comparative analysis have been made publicly accessible through]an interactive web tool called the <u>“Comparative Transcriptomic Placental Model Atlas (CTPMA)”</u>. This dataset is comprised of the normalized data that was summarized and cleaned together to remove all batch effects (N=12,793). We also include the 7,259 genes that were filtered out of our original dataset due to low and/or inconsistent expression, so that users have a comprehensive summary of the levels of expression of all genes, including those that may not be consistently expressed. This web tool includes several modules that will allow users to query this dataset. Specifically, users can search a given gene and identify the average gene expression data across all data and cluster memberships. We include a boxplot for each gene, across all model types, which will allow users to easily evaluate levels of expression for individual genes.

## Discussion

Human placental tissue is an important resource for studying the in-utero environment, however, use of primary tissue is limited by amenability to experimental methods and resources needed to recruit participants and collect tissue. We performed a transcriptome-wide comparison of gene expression of popular in vitro models of the human placenta, including 4 commercial cell lines as well as two in vitro models derived from primary tissue and human placental tissue from the CANDLE and GAPPS cohorts to inform selection of in vitro models for future experimental studies. We found that all 6 of the evaluated in vitro models had over 9,000 genes that were differentially expressed when compared to placental tissue samples, even with a conservative significance threshold of FDR<0.01. Villous explants, a primary tissue-based model, had the lowest number of DEGs (9,885), but overall, the large number of DEGs for each in vitro placental model, indicate that none of these common in vitro placental models are a close approximation of gene expression found in bulk placental tissue. This reinforces that in vitro model selection should be done based on expression of genes related to specific endpoints of interest for individual experiments and may vary based on the biological question.

The large number of DEGs (9,885) between the villous explants and bulk placental tissue was particularly surprising as the explants are derived from bulk placental tissue and would similarly contain a mixed population of maternal and fetal cells including fibroblasts, immune cells (Hofbauer, B cells, T cells, etc.), stem cells, and blood cells [39], rather than being strictly of trophoblast origin like most of the other models. The inclusion of immune cell types in the bulk placental tissue and villous explants is highlighted by the enrichment of the cytokine-cytokine receptor interaction KEGG pathway for Cluster 16 which has higher expression in both of these placental models. It is possible, however, that differences in sample processing or gestational age may be the cause of the transcriptomic differences between the villous explants and bulk placental tissue. The bulk placental tissue from CANDLE and GAPPS was frozen in RNALater after collection whereas, the villous explants from both studies were cultured fresh for 24-36 hours after collection [40,41]. Regarding gestational age differences, CANDLE and GAPPS bulk tissue was collected after delivery and therefore highlight gene expression at the end of gestation, whereas the villous explants were collected from terminated pregnancies in the 1^st^ [40] or 2^nd^ trimesters [41]. Bulk placental tissue from preterm (<37 week gestation) deliveries have previously been shown to exhibit transcriptomic differences from bulk placental tissue collected from term deliveries [23], so it is reasonable to expect there to be transcriptomic differences for the villous explants due to their collection at much earlier stages of placental development in the 1^st^ and 2^nd^ trimesters. While under ideal circumstances, we would compare bulk tissue and primary tissue based models collected during a similar time period of pregnancy, the scarcity of publicly available villous explant datasets precludes this.

Because some of the discrepancy in gene expression differences could be due to comparing bulk tissue gene expression to in vitro models that are largely from single cell types (with the exception of villous explants), we evaluated the differences between 4 commercial placental cell models and the primary trophoblasts against three primary tissue cell populations (EVTs, CTBs, and STBs) that were manually sorted by trophoblast cell type using FACS or villous tissue digests. Additionally, we evaluated the BeWo cell line in two stratified subsets of syncytialized and unsyncytialized cells to see if these designations matched their similarity to the sorted CTBs and STBs. Each of the cell models is considered to represent a different trophoblast cell type with HTR-8/SVneo representing EVTs, JAR, JEG-3, and unsyncytialized BeWo representing CTBs, and syncytialized BeWo and primary trophoblasts representing STBs. Such similarities provide support for our hypothesis that these cell lines would show the most similar gene expression profiles. This hypothesis did not hold true, as the comparisons to individual trophoblast cell types revealed that the primary trophoblast cell model was the most similar to all three trophoblast cell types (EVT, CTB, STB). This is likely related to its origin from a primary tissue source, while lack of immortalization may be more linked to gene expression similarities rather than the type of trophoblast cell represented by each model. Additionally, as JAR, JEG-3, and BeWo cells originated from gestational choriocarcinomas, rather than true placental tissue, it is not surprising that they do not closely approximate these trophoblast cell types derived from primary placental tissue.

Due to the extreme number of differentially expressed genes identified through our differential gene expression analysis, which limits the interpretability of those findings, we defined sets of genes that showed distinct expression profiles across the tested placental models using hierarchical clustering, which resulted in 53 clusters of genes. Pathway analysis of these 53 clusters identified 22 clusters with significantly enriched KEGG pathways that can be used to deduce which in vitro models are better-suited for the study of specific biological processes. For example, the Cytokine-Cytokine Receptor Interaction pathway was significantly enriched for genes within Clusters 13, 16, and 33. All three of these clusters show reduced gene expression in the choriocarcinoma cell lines (JAR, JEG-3, BeWo) compared to the other models indicating these models may not be appropriate for studying this pathway. After the placenta tissue, which has the highest average expression across these three clusters, the two in vitro models with the highest expression are villous explants and primary trophoblasts. While selecting the placental model with the highest expression may not be necessary for highly expressed genes, genes with lower expression levels in the placenta should be evaluated for sufficient expression levels in a model of interest before proceeding with experiments to ensure consistent and accurate results. We believe that evaluating the hierarchical clustering data for patterns of expression by placental model will be helpful for selecting an in vitro model that most closely recapitulates the gene expression of the human placenta for a particular study’s genes of interest.

As a proof of concept of this principal, we evaluated the clusters for membership and statistical overrepresentation of two sets of genes of interest including high-expression placental genes from the Human Protein Atlas (HPA) that may hold important roles in placental biology and ADME genes whose expression may be important for experiments evaluating toxicological or pharmaceutical agents. For example, three of the core ADME genes (*ABCB1, ABCC2, and ABCG2*) and 13 of the extended ADME genes are members of the ATP binding cassette family, made up of cell and organelle transmembrane proteins powered by ATP hydrolysis that are essential to a broad range of biological processes involved in the efflux transport of xenobiotics, hydrophobic drugs, metabolites, and polypeptides [42–44]. However, these 16 genes belong to 13 unique clusters with only Clusters 2, 6, and 17 containing at least two. Those hoping to study the role of these ATP binding cassette family genes or substrates that require their presence, would therefore need to carefully assess the expression profiles of 13 clusters to choose the placental model with the best expression profile for their study. In total, 6 of these genes (*ABCA4, ABCC2, ABCB6, ABCB8, ABCC9, ABCC11, ABCA1*) belong to clusters with expression that does not differ by placental model, however the remaining 10 ATP binding cassette family genes, belong to clusters with distinct expression profiles that differ by model, with primary trophoblasts being the model that has the most consistent expression across all 13 clusters.

Fetal sex is an important consideration when studying the placenta as there have been numerous studies that have illustrated differences in birth outcomes [45,46], placental physiology, and response to environmental insults that differ by fetal sex. Physiological differences in the placenta by fetal sex include differences in placental weight to birthweight ratio indicating potential differences in placental reserve capacity by fetal sex [47,48] as well as sex-specific transplacental signals to the brain [49]. Differences in placental gene expression by fetal sex have been noted during the first fetal androgen peak [50], in response to gestational exposures to environmental chemicals like phthalates and polycylic aromatic hydrocarbons [28–30], and at baseline due to epigenetic regulation of X- and Y-linked genes [51]. While fetal sex is typically reported for human placental tissue and primary tissue models, the fetal sex of commercial cell lines is not always readily available. To confirm fetal sex, we quantified gene expression on the Y-chromosome, which identified HTR-8/SVneo as the sole tested commercial cell line of female origin, while the three choriocarcinoma lines (JEG-3, JAR, and BeWo) were of male origin (Figure 6, Supplemental Table 4). Based on the results of this analysis, primary tissue-based models (Villous explants and primary trophoblasts), which typically have known fetal sex data, tended to show the most similar gene expression profiles to human placenta tissue. We believe that knowing the fetal sex of commercial cell lines is also important for contextualization and interpretation of results in light of physiological differences between male and female placentas that extend to differences in gene expression.

To our knowledge, this study represents the first comprehensive evaluation of the transcriptome of placental in vitro models that has directly made comparisons to the bulk placental transcriptome from human tissue samples. Strengths of this study include that it utilized publicly available RNA sequencing data and evaluated six common in vitro models of the placenta. However, the use of publicly available data was also a limitation as the control samples used in this analysis underwent various control treatments (i.e. DMSO, non-targeting siRNA, etc.), as well as that we have different sample numbers within each dataset and for each of the in vitro placental models. The use of different cell culture conditions (length of culture, cell media composition, and plated cell density) can also have an effect on gene expression [52]. To account for these differences, we adjusted for batch effects and unknown confounding using the method RUVr and visualized and quantified the variance in our data through principal components analysis which found that despite differences in sample size and treatment that the samples grouped by placental model. The inclusion of 1083 human placental tissue samples from the CANDLE and GAPPS cohorts is a significant strength of this work, which allowed us to directly compare the transcriptome of the in vitro models to that of bulk placental tissue. Future studies can expand upon our research by investigating the comparability of additional placental in vitro models that were not included in the present study such as the SWAN-71 [53] and ACH-3P cell lines [54], human trophoblast stem cells [55], or trophoblast organoids [56]. Inclusion of some of these models such as the human trophoblast stem cells and trophoblast organoids would be an important advancement for this work due to their growing utility as in vitro models of the placenta [57], but is hindered by the recency of their development, limited publicly available data, and the existence of subtypes of organoids (primary tissue or stem cell based) that are derived using separate protocols which have been shown at single-cell resolution to produce distinct transcriptomes [58]. Additional depth can also be added to this investigation through evaluation of genetic polymorphisms, RNA stability and protein translation in placental tissue and in vitro models.

To broaden the use of this data for future research, we have developed the <u>“Comparative</u> <u>Transcriptomic Placental Model Atlas (CTPMA</u>)”, a web tool that summarizes these results. This web application will empower research teams to evaluate their selected in vitro placental model for suitability during experiment planning and have a better understanding of the limitations of their selected model based on inherent gene expression differences between human placental tissue and their selected in vitro model of the human placenta.

In summary, our results indicate that there is no clear placental cell line that is a close approximation to human placental tissue. Researchers need to determine the best model system based on their individual research question, which may be inherently related to a specific biological pathway. In this manuscript and in the corresponding web application, we provide a global summary of gene-level and pathway level data for each in vitro placental model compared to human placental tissue, enabling researchers to better evaluate biological models. In the future, this tool can be improved through integration of multi-omics data, as well as integration of other in vitro placental models as they become available.

## Acknowledgements

This research was conducted using 1) specimens and data collected and stored on behalf of the Global Alliance to Prevent Prematurity and Stillbirth (GAPPS) Repository and 2) specimens and data collected from the Conditions Affecting Neurocognitive Development and Learning in Early Childhood (CANDLE) study. We would like to thank the CANDLE and GAPPS participants as well as the research staff and clinicians that facilitated these studies. We would also like to acknowledge the numerous studies that made their RNA sequencing data publicly available on GEO or SRA. We would also like to acknowledge Seattle Children’s Research Scientific Computing and Jenny Smith for their help with data management for this project. The content is solely the responsibility of the authors and does not necessarily represent the official views of the National Institutes of Health. This manuscript has been reviewed by PATHWAYS for scientific content and consistency of data interpretation with previous PATHWAYS publications.

## Supplemental Figures

**Supplemental Figure 1:**
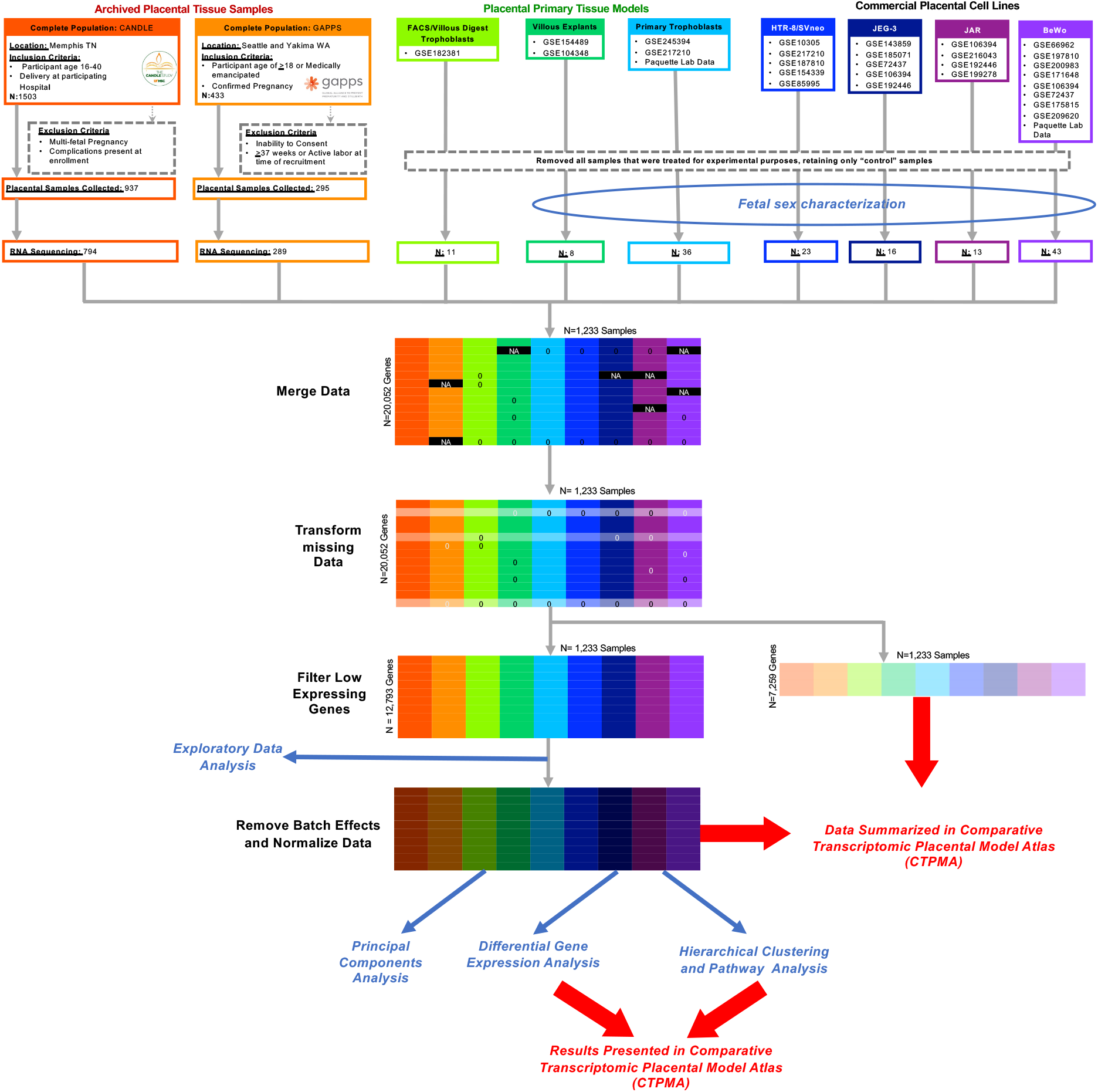
Methods diagram of data acquisition, cleaning, and analysis used for input to the <u>Comparative Transcriptomic Placental Model Atlas (CPTMA)</u>

